# Persistence of SARS-CoV-2 and its surrogate, bacteriophage Phi6, on surfaces and in water

**DOI:** 10.1101/2023.03.14.532590

**Authors:** Ana K. Pitol, Samiksha Venkatesan, Michael Hoptroff, Grant L. Hughes

## Abstract

The COVID-19 pandemic has motivated research on the persistence of infectious SARS-CoV-2 in environmental reservoirs such as surfaces and water. Viral persistence data has been collected for SARS-CoV-2 and its surrogates, including bacteriophage Phi6. Despite its wide use, no side-by-side comparisons between Phi6 and SARS-CoV-2 exist. Here, we quantified the persistence of SARS-CoV-2 and Phi6 on surfaces (plastic and metal) and in water and evaluated the influence that the deposition solution has on viral persistence by using four commonly used deposition solutions: two culture media (DMEM and Tryptone Soya Broth (TSB)), Phosphate Buffered Saline (PBS), and human saliva. Phi6 remained infectious in water significantly longer than SARS-CoV-2, having a half-life of 27 hours as compared with 15 hours for SARS-CoV-2. The persistence of viruses on surfaces was significantly influenced by the virus used and the deposition solution, but not by the surface material. Phi6 remained infectious significantly longer than SARS-CoV-2 when the inoculation solution was culture media (DMEM, TSB) and saliva. Using culture media and saliva led to half-lives between 9 hours and 2 weeks for Phi6, as compared to 0.5 to 2 hours for SARS-CoV-2. Using PBS as a deposition solution led to half-lives shorter than 4 hours for both viruses on all surfaces. Our results showed that, although it has been frequently used as a surrogate for coronaviruses, bacteriophage Phi6 is not an adequate surrogate for studies quantifying SARS-CoV-2 persistence, as it over-estimates infectiousness. Additionally, our findings reveal the need of using adequate deposition solutions when evaluating viral persistence on surfaces.

## INTRODUCTION

SARS-CoV-2 is primarily transmitted through respiratory droplets and aerosols^1^. Nevertheless, the RNA of SARS-CoV-2 has been extensively detected on environmental reservoirs, with concentrations as high as 10^5^ genome copies (gc) per swab on surfaces, and >10^5^ gc mL^-1^ in wastewater samples^2–7^. The extensive contamination of environmental reservoirs with the SARS-CoV-2 RNA has been the source of concern throughout the COVID-19 pandemic^8^. However, viral RNA concentration is higher than that of infectious viral particles on a given sample^9,10^. Hence, using data on RNA contamination to estimate the risks associated with people interacting with contaminated environments could lead to overestimating the risks. Therefore, it is important to understand the persistence of infectious SARS-CoV-2 to estimate the magnitude of the risks associated with people interacting with contaminated environments.

Since the beginning of the pandemic, there have been a number of studies quantifying the persistence and inactivation of SARS-CoV-2 on surfaces and in liquids under different environmental conditions such as temperature and humidity^9–16^. However, there is a lack of consistency in the experimental designs of these studies, which could lead to biased outcomes. For example, in studies of SARS-CoV-2 persistence on surfaces, a wide range of deposition solutions has been used to inoculate the virus, these range from culture media and Bovine Serum Albumin (BSA) containing media^16,17^ to bodily fluids such as human saliva and mucus^10,18^. The solutions used for virus inoculation have very different characteristics which can influence viral survival^19^. Additionally, working with SARS-CoV-2 requires handling the virus in high containment facilities, which presents many challenges and limits the type of experiments that can be performed. For example, it is not advisable to do experimental work using SARS-CoV-2 with human volunteers, such as testing SARS-CoV-2 persistence on human hands or quantifying the transfer of the virus between surfaces and hands.

Therefore, efforts have been made to use surrogates, instead of SARS-CoV-2, to understand the mechanisms of its survival, inactivation, and transfer ^20–24^. Bacteriophages are frequently used as surrogates of pathogenic viruses because they are safe, inexpensive, and do not require contaminant facilities ^25–27^. Bacteriophage Phi6 is one of the few bacteriophages that has a lipid envelope ^28^. Therefore, it has been used as a surrogate for enveloped viruses such as SARS and MERS coronaviruses, influenza virus, and Ebola ^20–22,27,29–32^. To this end, bacteriophage Phi6 has been used as a surrogate for SARS-CoV-2 in studies evaluating virus persistence on surfaces ^22,24,32^, virus persistence in water and wastewater ^33^, virus inactivation ^20,21,30^, virus transfer from surfaces to hands^34^ and virus recovery from fingertips ^23^. While these studies provide important information on viral persistence and transfer, there is a gap in our knowledge related to the adequacy of using bacteriophage Phi6 as a proxy for SARS-CoV-2. As such, there is an urgent need to evaluate the suitability of using bacteriophage Phi6 as a surrogate for SARS-CoV-2 in persistence studies.

Here we evaluated the persistence of SARS-CoV-2 and bacteriophage Phi6 in water. Additionally, we assessed the persistence of SARS-CoV-2 and Phi6 on surfaces of two different non-porous materials: plastic (PVC) and metal (stainless steel). Since the deposition solution used to inoculate the virus on the surface could have an influence on virus persistence, we evaluated four commonly used deposition solutions: Phosphate Buffered Saline (PBS), Tryptic Soy Broth (TSB), cell culture media (DMEM), and human saliva. Finally, we used the data to assess the suitability of using bacteriophage Phi6 as a surrogate for SARS-CoV-2 in studies of virus persistence in water and on surfaces.

## METHODS

### Bacteriophage Phi6 production and enumeration

Bacteriophage Phi6 (DSM 21518) and its host *Pseudomonas syringae* (DSM 21482) were obtained from the DSMZ (Deutsche Sammlung von Mikroorganismen und Zellkulturen). To culture the bacteriophage Phi6, we used a protocol adapted from Pitol *et al.* 2017 ^35^. Briefly, 100 mL of Tryptic Soy Broth (TSB; Millipore) containing log-phase *P. syringae* was inoculated with 100 μL of a 10^8^ Plaque Forming Units (PFU) mL^-1^ stock of bacteriophage Phi6 and incubated overnight. The following day, the media was centrifuged for 15 min at 5000 rpm and the supernatant was filtered using a 0.45 μm filter unit. Aliquots of the supernatant with a final concentration of 1.5 x10^11^ PFU mL^-1^ were stored at 4°C for subsequent assays.

To enumerate Phi6 we used the standard Double Agar Layer assay^35^. Briefly, soft TSB agar (0.5% Agar) was inoculated with 100 μL of an overnight culture of *P. syringae* and 100 μL of the sample containing Phi6. Samples were mixed and poured on hard agar (1.5%. Agar) TSB plates and incubated at 25°C overnight. Negative controls (100 μL of an overnight culture of *P. syringae* with no bacteriophage) were included in each experiment. All dilutions were quantified in duplicates, and all experiments were performed in triplicates.

### SARS-CoV-2 production and enumeration

SARS-CoV-2, Delta variant, passage 4 (SARS-CoV-2/human/GBR/Liv_273/2021, OK392641)^36^ was amplified and quantified using Vero E6 cells (African green monkey kidney cells, Public Health England). Vero cells were maintained at 37°C and 5% CO_2_ in Dulbecco’s Minimal Essential Medium (DMEM; Corning) supplemented with 10 % Foetal Bovine Serum (FBS; Sigma-Aldrich) and 0.05 mg mL^-1^ of gentamicin (Gibco). To amplify SARS-CoV-2, a flask T-150 of confluent Vero E6 was inoculated with 20 μL of a 10^6^ PFU mL^-1^ stock of SARS-CoV-2 Delta variant, passage 4, and incubated for 72h. Subsequently, the media was recovered and centrifuged for 15 min at 5000 rpm to remove the remaining cells and cell debris. Finally, the stock solution, SARS-CoV-2 passage 5 at a concentration of 10^7^ PFU mL^-1^, was concentrated using the Amicon Ultra Centrifugal Filter (100kDa; Merk Millipore Amicon) to a final concentration of ~10^8^ PFU mL^-1^.

Standard plaque assay was performed to quantify infectious virus as previously described ^37^. Briefly, samples were serially diluted and inoculated in a confluent monolayer of Vero E6 cells. One hour after infection, an agarose media overlay (2% Agarose in DMEM supplemented with 2% FBS) was applied to the cell monolayer and incubated at 37°C and 5% CO_2_ for 72h. Subsequently, the cells were fixed with Formalin 10% (VWR International) and stained with Crystal Violet (Sigma-Aldrich) and plaques were counted. All experiments were conducted in a containment level 3 laboratory by personnel trained in the relevant code of practices and standard operating procedures.

### Surfaces and deposition solutions

To study the persistence of SARS-CoV-2 and Phi6 on surfaces, each virus was suspended in four different deposition solutions to a final concentration of ~10^6^ PFU mL^-1^. The solutions used were: Phosphate Buffered Saline (PBS; Gibco), Tryptic Soy Broth (TSB; Millipore), DMEM supplemented with 2 % FBS, and human saliva. Saliva was collected at Unilever Research Port Sunlight as described elsewhere ^36,38^. Briefly, healthy donors provided a stimulated daytime saliva sample for which they were given a piece of gum to chew (Wrigley’s Turbulence). Subjects were given a maximum of five sterile 30 ml containers in which they were asked to provide a 20–25 ml sample of saliva per container. Subjects were requested to leave 30 min after eating or drinking before providing a saliva sample. Saliva samples were stored overnight at −80°C prior sterilization using to gamma irradiation (Systagenix, UK, Cobolt 60 turntable, dose rate 1.2 kGy/h, minimum dose 32.1 kGy). After sterilization, saliva was stored at 4°C until used.

To study virus persistence on surfaces, we selected two non-porous materials: Plastic (PVC plastic vinyl), and metal (stainless steel). Circular coupons of stainless steel and plastic were disinfected by soaking in 70% ethanol (VWR International) for 30 minutes. Subsequently, the coupons were thoroughly rinsed with deionized water before allowing them to dry inside a class II biological safety cabinet for one hour.

### Persistence of SARS-CoV-2 and Phi6 on surfaces

The plastic and steel coupons were inoculated by pipetting a 1 μL droplet of deposition solution (PBS, TSB, DMEM, saliva) containing ~10^6^ PFU mL^-1^ of virus (SARS-CoV-2 or Phi6) in the centre of the coupon. The inoculated coupons were incubated at 25°C in a container partially opened to allow evaporation of the droplets. Coupons were sampled at different time points by pipetting up and down 15 times using 50 μL cell culture media (DMEM supplemented with 2% FBS) and samples were stored at −80°C before being quantified together at the end of the experiment. Positive controls, consisting of 50 μL DMEM inoculated with 1 μL of virus (SARS-CoV-2 or Phi6) suspended in each of the four viral matrices (PBS, TSB, DMEM, Saliva), were run alongside each treatment. Experiments were repeated three times.

### Persistence of SARS-CoV-2 and Phi6 in water

A stock of 10^8^ PFU mL^-1^ of virus (SARS-CoV-2 or Phi6) was serially diluted in mineral water (Volvic Natural Mineral Water) to a final concentration of 10^6^ PFU mL^-1^ and aliquoted in samples of 50 μL. The samples were incubated at 25°C and recovered at different time points. Subsequently, they were diluted in 200 μL of culture media (DMEM supplemented by 2% FBS) to a final volume of 250 μL, and stored at - 80°C, before being quantified. Positive controls, consisting of 200 μL DMEM inoculated with 50 μL of virus (SARS-CoV-2 or Phi6) suspended in water, were collected alongside each treatment. Experiments were repeated three times.

### Data analysis

All data analysis was performed using R statistical software (Version 4.1.0). Multiple regression analysis was used to assess the difference in the number of viruses on the water as a function of time and virus (Phi6, SARS-CoV-2). In the experiments which determined virus persistence on surfaces, multiple regression analysis was used to evaluate the survival of viruses as a function of virus, deposition solution (PBS, TSB, DMEM, saliva), surface material (plastic, steel), and time.

Linear regression was used to calculate the first-order decay constant, *k* (h^-1^), as the slope of ln (*N*/*N*_0_) versus time in hours (Equation 1), where *N* is the number of viruses at time = t, and *N*_0_ is the number of viruses at the beginning of the experiment (t=0). Data with concentrations below the limit of detection (LOD) was excluded from the regression analysis. The decay constant, *k,* was used to estimate the half-life of the virus, *t*_50_ (h^-1^), and the time required for 90% of the reduction in the number of viruses, *t*_90_ (h^-1^). The *t*_50_ and the *t*_90_ were estimated using equations 2 and 3 respectively.

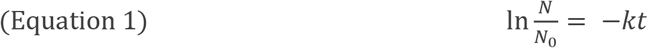

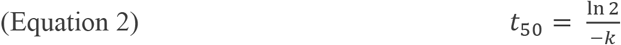

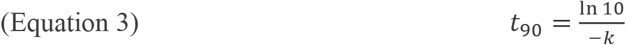

## RESULTS AND DISCUSSION

### SARS-CoV-2 and Phi6 survival in water

A multiple linear regression (MLR) was calculated to predict the concentration of viruses in the water as a function of time and virus type (F (2,59) = 29.73, p<0.001), R2=0.49). The persistence of the viruses in water was significantly influenced by the virus used (MLR, p<0.001). The half-life of bacteriophage Phi6 was 27 hours as compared with 15 hours for SARS-CoV-2 (Table 2, Figure 2). Therefore, bacteriophage Phi6 significantly overestimated the persistence of SARS-CoV-2 in the water. The decay rates reported here for both viruses, Phi6 and SARS-CoV-2, fall within the range of data reported elsewhere^33,39^. The time taken to achieve a 90% reduction in the titter (*t*_90_) for SARS-CoV-2 in mineral water was 50 hours. Our findings were comparable to other studies using autoclaved or filtered river water and tap water at similar temperatures (20-25°C), *t*_90_= 48-79.2 hours ^14,15,40^. Additionally, we obtained a *t*_90_ of 89 hours for Phi6, which is consistent with previous research that shows a range of *t*_90_ between 74 and 179 hours for autoclaved or filtered river water and tap water at similar temperatures ^27,33^. The variation observed in virus survival between different studies can be explained by several factors including the chemical composition of the water used in the experiments (e.g., oxygen level, pH, presence of organic matter) and environmental variables such as light exposure and temperature ^41^. Therefore, a quantitative comparison between different studies is difficult, which highlights the importance of performing a side-by-side comparison between the viruses when evaluating the suitability of using viral surrogates.

**Figure 1.**
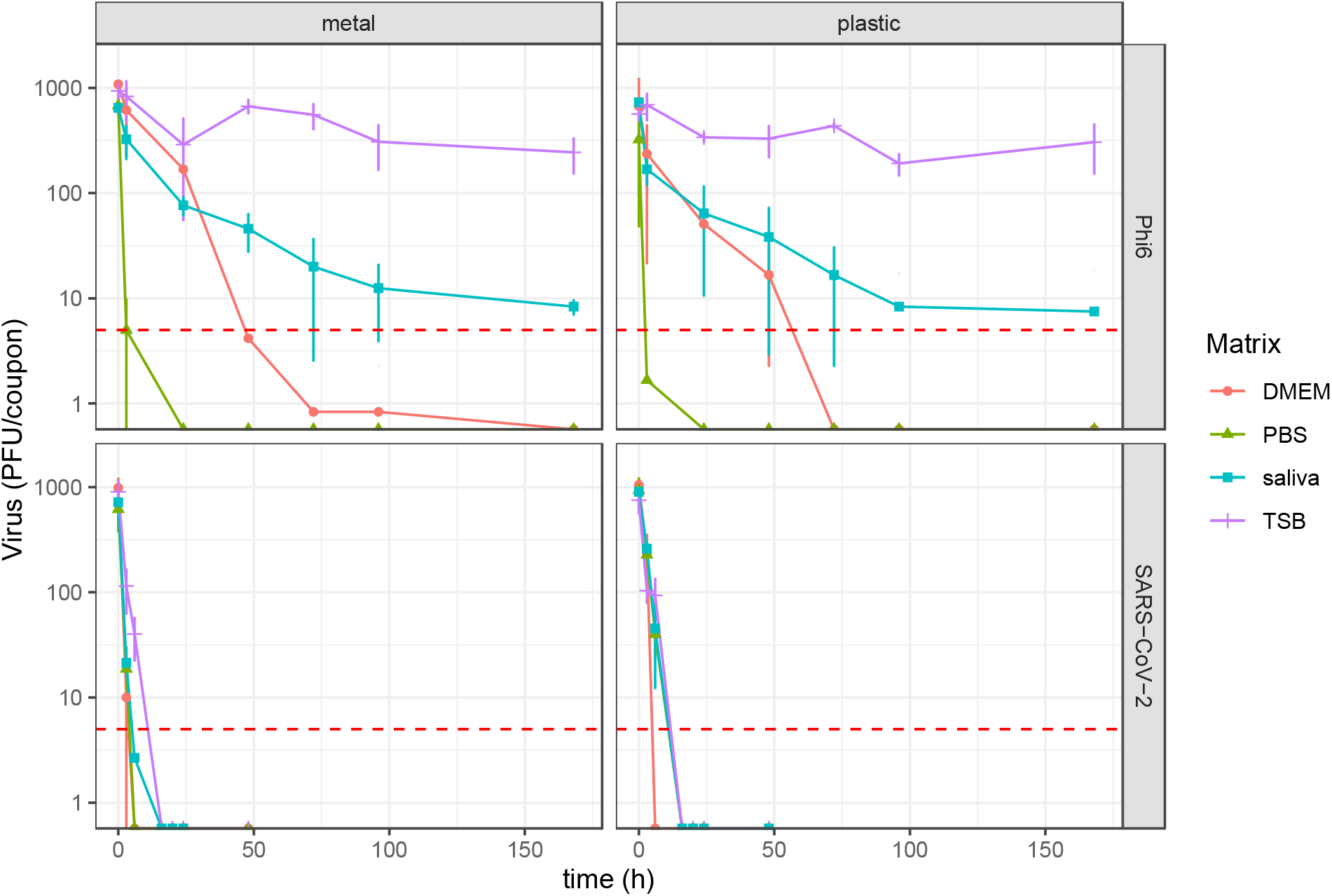
Virus survival on surfaces as a function of surface material (plastic vs metal), virus type (SARS-CoV-2 vs Phi6) and deposition solution (DMEM, PBS, TSB, and saliva). Data shows the mean and the standard deviation of triplicates. Red dotted line shows the limit of detection of the assays.

**Figure 2.**
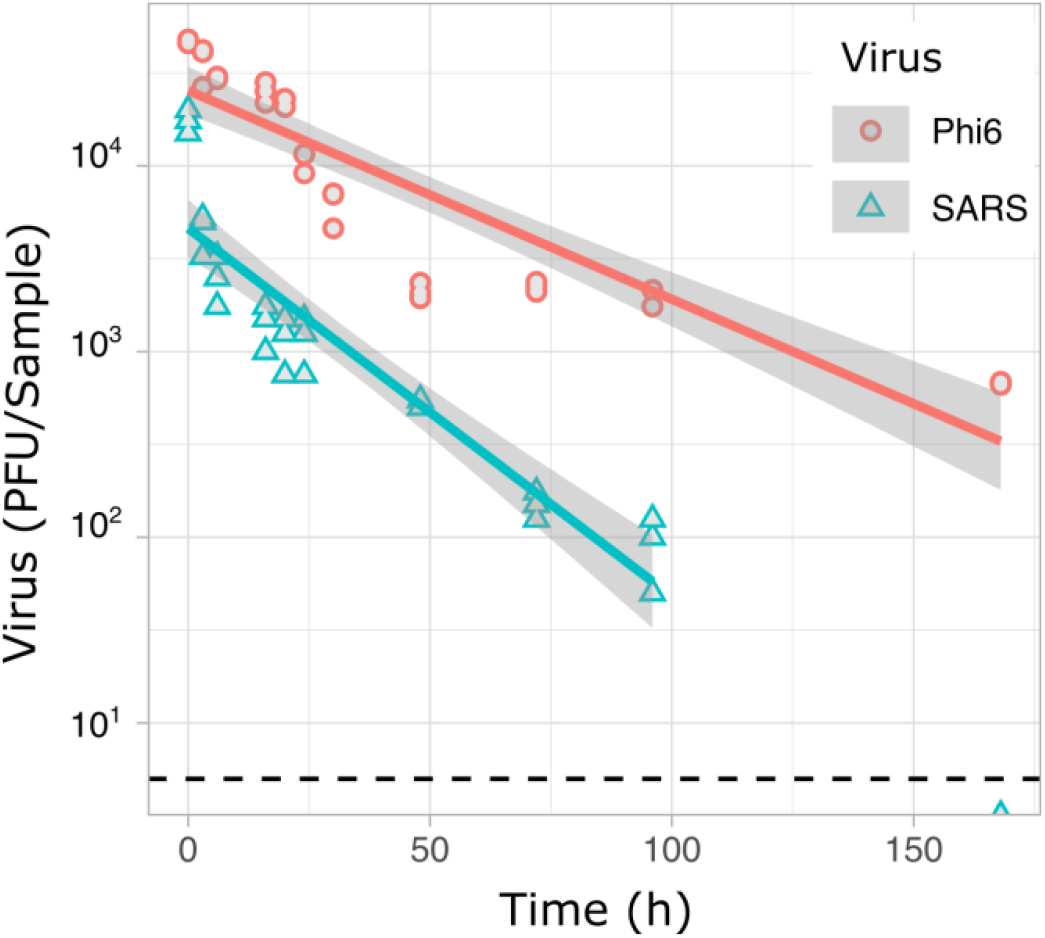
Persistence of SARS-CoV-2 and bacteriophage Phi6 on mineral water. Number of virus per sample are shown in red circles for Phi6 and blue triangles for SARS-CoV-2. The red and blue continuous lines show the fitted linear model for Phi6 and SARS-CoV-2 respectively, with grey areas showing the 95% CI for each regression line. The black dotted line shows the limit of detection for the assays.

### Persistence of SARS-CoV-2 and Phi6 on surfaces

MLR was used to predict the concentration of viruses as a function of time, virus type, deposition solution, and surface material (F (6, 376) = 12.01, p<0.001). Virus type, deposition solution, and time were significant predictors of virus concentration (MLR, virus type p<0.001, deposition solution p<0.001, time p<0.001). Conversely, the influence of surface material (metal vs plastic) on virus persistence was not statistically significant (p=0.57). The most marked observation to emerge from the data comparing the survival of SARS-CoV-2 with that of bacteriophage Phi6 on surfaces was that Phi6 survives significantly longer than SARS-CoV-2 (Figure 1, Table 1). The half-life of Phi6 was 403 hours (~2 weeks) on plastic and 250 hours (~10 days) on metal when the deposition solution was TSB. Conversely, inoculating the surfaces with SARS-CoV-2 suspended in TSB resulted in half-lives of 1.3 and 1.9 hours for metal and plastic respectively. Similar patterns of higher persistence of Phi6 as compared with SARS-CoV-2 were obtained when using saliva and DMEM as deposition solutions (Table 1). The half-life of Phi6 on plastic was 1-3 days when the deposition solution was saliva, as compared to 0.8-1.2 hours for SARS-CoV-2. Using DMEM as deposition solution, the half-life of Phi6 on plastic and metal were 12.3 and 9.3 hours respectively, as compared with halve-lives of 1.3 and 0.5 hours for SARS-CoV-2. Interestingly, when viruses were suspended in PBS, their half-lives were short, regardless of the virus used. Using PBS as a deposition solution led to Phi6 half-lives of 1.7 hours and 1.5 hours in plastic and metal respectively, as compared to 4 and 2 hours for SARS-CoV-2.

**Table 1.** Summary of the Mean Linear Regression Model Parameters for the Persistence of SARS-CoV-2 and Phi6 on Surfaces.

| Virus | Deposition solution | Surface material | T<sub>90</sub> (h) | T<sub>50</sub> (h) | k, SE k (h<sup>-1</sup>) |
| --- | --- | --- | --- | --- | --- |
| SARS-CoV-2 | DMEM | plastic | 4.24 | 1.28 | -0.543, 0.125 * |
| SARS-CoV-2 | TSB | plastic | 6.35 | 1.91 | -0.363, 0.080 |
| SARS-CoV-2 | PBS | plastic | 4.27 | 1.29 | -0.539, 0.042 |
| SARS-CoV-2 | Saliva | plastic | 4.13 | 1.24 | -0.557, 0.099 |
| SARS-CoV-2 | DMEM | metal | 1.63 | 0.49 | -1.414, 0.086 * |
| SARS-CoV-2 | TSB | metal | 4.32 | 1.30 | -0.533, 0.063 |
| SARS-CoV-2 | PBS | metal | 2.00 | 0.60 | -1.150, 0.223* |
| SARS-CoV-2 | Saliva | metal | 2.68 | 0.81 | -0.859, 0.138 |
| Phi6 | DMEM | plastic | 40.81 | 12.28 | -0.056, 0.031 |
| Phi6 | TSB | plastic | 1338.77 | 403.01 | -0.002, 0.001 |
| Phi6 | PBS | plastic | 1.66 | 0.50 | -1.389, 0.000* |
| Phi6 | Saliva | plastic | 268.96 | 80.96 | -0.009, 0.002 |
| Phi6 | DMEM | metal | 31.03 | 9.34 | -0.074, 0.009 |
| Phi6 | TSB | metal | 829.92 | 249.83 | -0.003, 0.001 |
| Phi6 | PBS | metal | 1.51 | 0.46 | -1.520, 0.086 |
| Phi6 | Saliva | metal | 93.92 | 28.27 | -0.025, 0.003 |

\*Only 2 time points were considered in the regression, as the virus was inactivated within 6 hours.

**Table 2.** Linear Regression Models for Persistence of SARS-CoV-2 and Bacteriophage Phi6 on Water

| Virus | Liquid Matrix | $t_{90}$<br>(h) | $t_{50}$<br>(h) | $k$, SE $k$<br>(h<sup>-1</sup>) |
| --- | --- | --- | --- | --- |
| SARS-CoV-2 | Mineral water | 50.44 | 15.19 | -0.046, 0.004 |
| Phi6 | Mineral water | 88.93 | 26.77 | -0.026, 0.002 |

$t_{90}$ : Time required for 90% of the reduction in the number of viruses (hours)
$t_{50}$ : Half-life or time required for 50% of the reduction in the number of viruses (hours)
$k$ : First order decay constant (hours<sup>-1</sup>)

In addition to the higher persistence of bacteriophage Phi6 over SARS-CoV-2, our data indicate that the deposition solution had a significant influence on virus survival (Table 1, Figure 1). This finding is consistent with previous research that suggest that the survival of viruses, including SARS-CoV-2, is highly influenced by the liquid matrix used to suspend the virus^19,42^. Pastorino *et al.* showed that higher protein content in the deposition solution increases viral persistence on surfaces^19^. In the present study we used four deposition solutions: TSB (20 grams of protein per litter, g L^-1^), human saliva (0.7 - 2.5 g L^-1^, based on data by Lin *et al*^43^), DMEM supplemented with 2% FBS (~1.2 g L^-1^), and PBS (0 g L^-1^). Our data showed that bacteriophage Phi6 survives the longest in TSB, followed by saliva, DMEM, and lastly PBS, which correlates with protein concentration of the deposition solution. Other components within deposition solutions that have been shown to influence the stability of the viruses is the pH ^42,44^, salt concentration ^42,45^, and the presence of polysaccharides ^46^ in the solution.

Our data on SARS-CoV-2 survival on surfaces are consistent with multiple research studies that showed that the half-life of SARS-CoV-2 at 20-27°C was between 1.5 and 9 hours in plastic surfaces ^10,13,16,47^, and between 3.4 and 7.8 hours in stainless steel ^9,13,16^. Nevertheless, other studies have shown a longer survival rate for SARS-CoV-2 on surfaces at similar temperatures, with half-lives in the order of days rather than hours ^17,48^. Differences between the studies include using different deposition solutions, inoculum volumes, light exposure during the experiment, and drying times. Since factors such as temperature, light exposure, and experimental variables such as the composition of the deposition solution used to inoculate the surfaces play a crucial role in the survival of viruses, it is desirable for future research to carefully select these factors when evaluating the survival of emerging pathogens in the environment.

We acknowledge our study has limitations, which are inherent when working with a category 3 pathogen such as SARS-CoV-2. For example, due to biosafety requirements, samples were placed in a container inside an incubator with no light or active air movement, which are factors that have been shown to influence virus survival^41,49^. Therefore, the decay of the viruses presented here may not adequately reflect the decay that viruses have in scenarios with sunlight exposure and air movement. Additionally, experiments were performed at ambient humidity, which fluctuated between 25 and 50%. Temperature and humidity have shown to have an influence on the persistence of SARS-CoV-2 on the surfaces ^13,18^. For example, one study showed that the half-life of SARS-CoV-2 in stainless steel ranges from 3 to 70 hours, depending on the environmental temperature and humidity ^13^. Therefore, adequate control of both temperature and humidity is advisable to when evaluating the persistence of pathogens.

Despite the limitations of this study, our findings highlight the importance of evaluating the suitability of using viral surrogates by performing side-by-side comparisons with the pathogen of interest to control all variables that could potentially influence the outcome. Our results showed that, although it has been frequently used as a surrogate for coronaviruses, bacteriophage Phi6 is not an adequate surrogate for studies quantifying SARS-CoV-2 persistence, as it over-estimates infectiousness. Additionally, our findings reveal the need of using adequate deposition solutions when evaluating viral persistence on surfaces. Future research on the persistence of pathogens should place careful consideration on the methodology used, selecting a deposition solution, inoculation method, and environmental parameters that adequately mimic real-life scenarios.

## ACKNOWLEDGMENT

Carole Philpotts (Unilever) is thanked for her assistance in supply of saliva. This work was supported by UKRI grants (20197 and 85336) awarded to GLH. GLH was further supported by the BBSRC (BB/T001240/1, BB/V011278/1, and BB/W018446/1), the UKRI (85336), the EPSRC (V043811/1), a Royal Society Wolfson Fellowship (RSWF\R1\180013), the NIHR (NIHR2000907) and the Bill and Melinda Gates Foundation (INV-048598). This work was additionally supported by a Pandemic Institute grant (TPIMPX01) awarded to AKP. The Pandemic Institute is formed of seven founding partners: The University of Liverpool, Liverpool School of Tropical Medicine, Liverpool John Moores University, Liverpool City Council, Liverpool City Region Combined Authority, Liverpool University Hospital Foundation Trust, and Knowledge Quarter Liverpool. The views expressed are those of the author(s) and not necessarily those of the Pandemic Institute.

